# Antioxidant Polysulfide Nanoparticles Ameliorate Ischemia-Reperfusion Injury and Improve Porcine Kidney Function Post-Transplantation

**DOI:** 10.1101/2023.06.21.545864

**Authors:** John P Stone, Richard D’Arcy, Abbey Geraghty, Kavit R Amin, Matilde Ghibaudi, Nora Francini, Angeles Montero-Fernandez, Dilan Dabare, Giulia Coradello, Jo Bramhall, Nicholas Galwey, Marieta Ruseva, Nicola Tirelli, James E Fildes

**Author notes:** **Correspondence to:** Prof James E Fildes.

## Abstract

Ischemia-reperfusion injury (IRI) is a significant complication in kidney transplantation, often affecting the viability and function of organs. Normothermic machine perfusion (NMP) is a technique used to improve the condition of organs prior to transplantation. In this study, we show that incorporating antioxidant poly(propylene sulfide) nanoparticles (PPS-NPs) during cold-storage and NMP significantly enhances its efficacy in reducing IRI upon porcine kidney transplantation. We found that by scavenging reactive oxygen species, PPS-NPs reduced oxidative stress and inflammation that occurs during ischemia-reperfusion with oxidized DNA reduced 5.3x and both TNF-α and complement activation approximately halved. Our studies show that this approach led to significantly improved hemodynamics, better renal function, and tissue health compared to NMP alone. The results suggest that incorporating PPS-NPs into transplantation protocols may expand the pool of kidneys suitable for transplantation and enhance overall transplantation success rates. The broader impact of this work could extend to other organ transplants, suggesting a wider application for nanoantioxidant technologies in organ preservation.

## Introduction

Kidney transplantation remains the mainstay treatment option for end stage renal disease, yet there is a critical mismatch between the high demand for transplants and the limited supply of available organs, posing a significant public health challenge. In the United States alone, approximately 100,000 patients await a kidney transplant, yet only 25,000 transplantations are performed each year^1^; alarmingly, 6,000 individuals die each year while waiting for a transplant^2^. Despite the urgent need, a staggering 25% of deceased donor kidneys (∼5000) were discarded in 2022.^3^ This loss of potentially viable organs has drawn national attention, leading to the launch of the Advancing American Kidney Health initiative by the White House, coupled with concerted efforts by advocacy groups like the National Kidney Foundation to minimize organ waste and enhance transplant accessibility.^4^ These initiatives highlight the urgent need for innovative techniques and therapies that can effectively increase the viability of available donor kidneys and reduce the tragic waste of life-saving resources.

The main determining factor in a kidney’s viability and ultimately transplant success, is the inevitability of ischemia reperfusion injury (IRI) (**Scheme 1**). This involves a cascade of events that begins at the point of organ retrieval and is further exacerbated at each stage of the transplant pathway. The current gold standard preservation technique of the donor kidney remains static cold storage (SCS), during which time there is no oxygen or nutritional support. As a result, metabolism becomes anaerobic, ATP is depleted, and there is a concurrent increase in intracellular calcium concentration and ion pump dysfunction. At the same time, pH decreases, resulting in tissue acidosis.^1^ Upon reperfusion, reactive oxygen species (ROS) such as superoxide or hydrogen peroxide increase due to the machinery for the conversion of molecular oxygen into ROS being upregulated under hypoxic conditions. Elevated ROS levels activate multiple immune pathways, including the complement cascade.^1^ Of note, ROS-mediated signaling may also contribute to maintaining active inflammatory pathways (through REDOX-sensitive transcription factors) with various origins.^2^

**Scheme 1.**
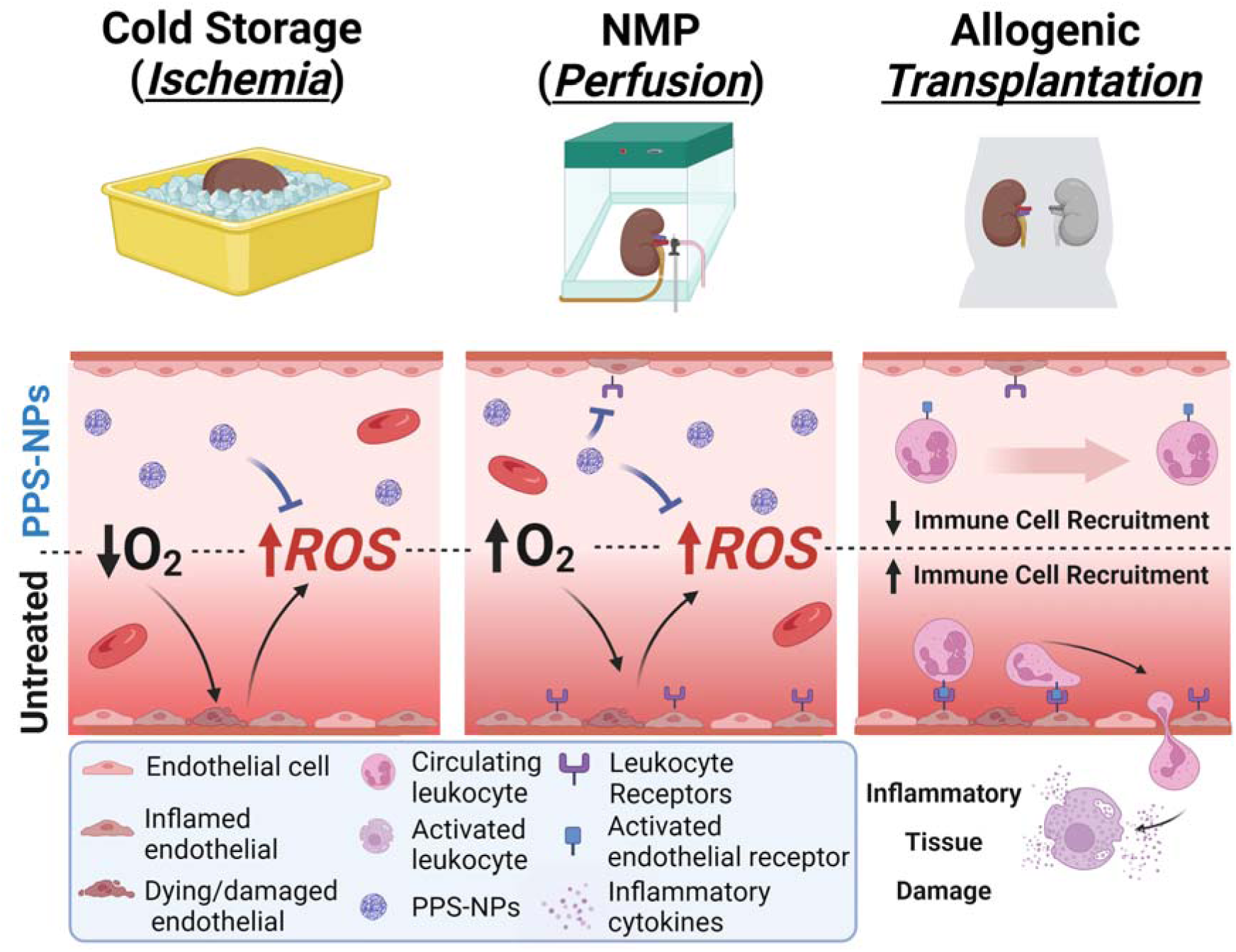
**Cold storage:** organs from deceased donors are typically stored on ice resulting in oxygen deprivation and increased ROS. This leads to endothelial cell damage and inflammation; inflamed endothelial cells present leukocyte receptors such as weak binding/rolling receptors (e.g., e-selectin) and ‘firm’ binding receptors such as ICAM-1 and VCAM-1. **Normothermic Machine Perfusion (NMP):** rewarming and reoxygenation during perfusion leads to a subsequent production of ROS and further endothelial damage and inflammation. **Transplantation:** native leukocytes from the transplant recipient bind to their receptors on inflamed endothelial cells in the transplanted organ resulting in extravasation and inflammatory activation/cytokine release – ultimately lowering organ viability and reducing transplant success odds.

Efforts to minimize IRI include improved cold storage strategies^5^, xenogeneic extracorporeal preservation^6–8^, and normothermic machine perfusion (NMP)^9^. We have worked towards the advancement of NMP^10–12^ (Scheme 2), which allows for the preservation of organs outside the body by maintaining them at normal body temperature while providing continuous oxygen and nutritional support. This technology enables 1) real time (ex vivo) evaluation and 2) the restoration of organs that might otherwise be considered marginal.^13^ Therapeutic intervention during NMP adds to a new and growing paradigm of ex vivo organ treatment which has the advantage of avoiding off-target side-effects seen with systemic delivery post-transplant.^14–17^ Whilst this approach to donor organ treatment is still nascent, nanoparticles (NPs) targeted to platelet-endothelial cell adhesion molecule-1 (CD31) or ICAM-2 demonstrate strong retention within kidneys during NMP with it hypothesized that these could potentially form slow-release, kidney localized drug depots after transplantation.^14, 17, 18^

**Scheme 2.**
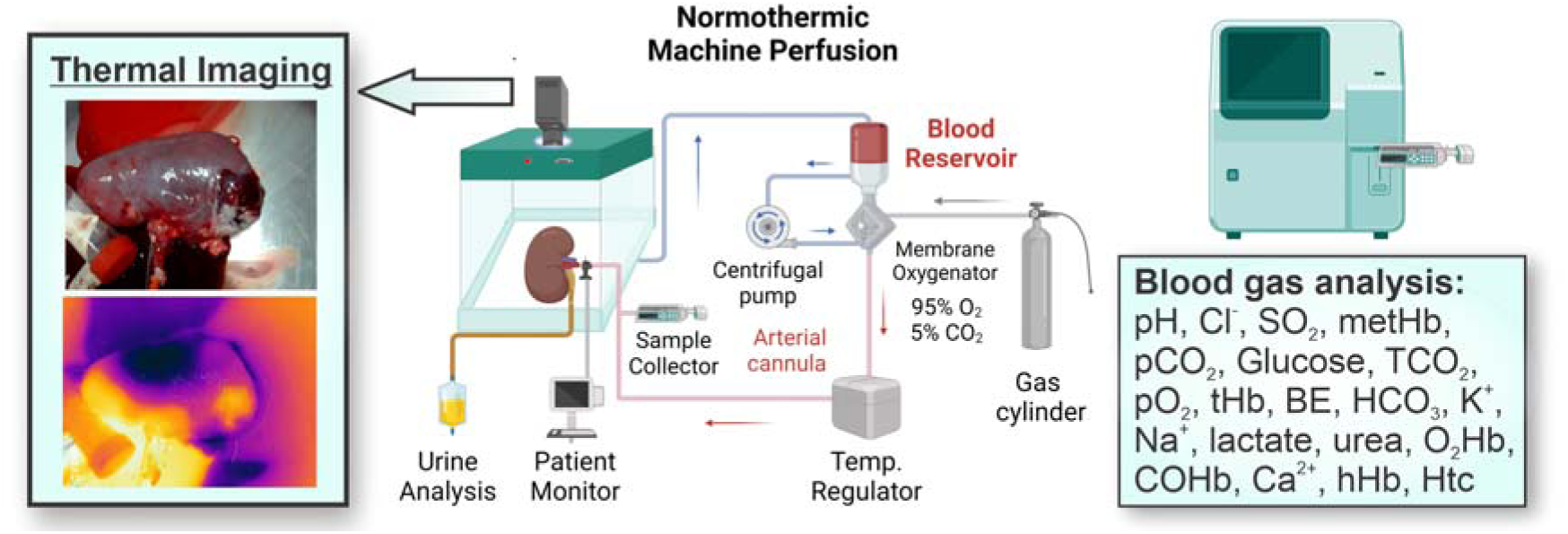
Detailed schematic of our NMP equipment setup used in this study.

Identified as a potential drug target, complement activation during both cold storage and upon transplantation significantly contributes to tissue injury and delayed graft function (DGF) following transplantation.^19^ Although various strategies aimed at directly inhibiting complement activation have been attempted, their success has been limited.^4^ Nevertheless, targeting the early initiation events of complement activation through ROS scavenging within the graft prior to transplantation may offer significant benefits. For instance, in one innovative approach to enhance transplant outcomes, anti-CD31 antibodies fused to the antioxidant enzyme catalase was administered to donor rats before lung retrieval. This targeted ROS scavenging not only mitigated IRI but also improved the function of the lung after transplantation.^5^

Remarkably, administering the therapy immediately after transplantation rather than prior to organ removal in the donor conferred no benefits. This lack of effect was attributed to substantial oxidative stress occurring during the cold storage phase prior to implantation, further highlighting the need to block ROS prior to transplantation. Inspired by this work, we sought to implement an antioxidant strategy during both SCS and NMP to improve transplant outcomes.

As an antioxidant, while catalase exhibits promising therapeutic properties, synthetic materials offer distinct translational advantages, including a broader spectrum of ROS quenching and a lower risk of immunogenicity. Additionally, synthetic antioxidant materials are not susceptible to denaturation or proteolytic degradation and when e.g., PEGylated, can achieve extended circulation half-lives. Antioxidant nanomaterials constructed from high-sulfur content organic polymers, such as poly(propylene sulfide) (PPS), offer a robust platform for scavenging ROS. These materials efficiently neutralize ROS through the sacrificial oxidation of their sulfur(II) atoms and have proven highly effective in managing oxidative stress. In PPS, this reaction predominantly targets hydrogen peroxide, producing an antioxidant effect analogous to that of catalase^20, 21^, but also potently scavenges other physiological ROS including hypochlorite, hydroxyl radical and peroxynitrite.^22–24^ As a result, PPS-based microparticles^25^, NPs^26^, micelles^27^ and other self-assembled colloidal species^28^ including gel-forming ones^29^ have been shown to exert significant anti-inflammatory effects in both *in vitro* and *in vivo* models of e.g. osteoporosis^27^, wound healing^29^, ischemic stroke^25, 26^, or traumatic brain injury^25^. Herein we further confirm the validity of PPS-NPs in a porcine kidney transplantation model and present what we believe to be the first therapeutic example of nanomedicine augmented NMP.

## Materials and Methods

### Synthesis and Fluorescent Labelling of Poly(propylene sulfide) nanoparticles (PPS-NPs)

Briefly, under an argon flow, 12.8 g of Pluronic F-127 was dissolved into 200 mL of degassed MilliQ water within a 250 mL round-bottom flask attached to a Radley’s Tornado Overhead Stirring manifold. To ensure complete dissolution/dispersion, the solution was stirred at 500 rpm for 10 minutes. To this solution, a mixture containing 175 mg of 2,2’-(ethylenedioxy)diethanethiol (0.96 mmol) and 6.307 g propylene sulfide (86.4 mmol, 45 equiv.s per thiol) was then added. The stir speed was increased to 1,000 rpm and the solution stirred for a further 10 minutes. To initiate the polymerization, 292 mg of DBU (1.92 mmols, 1 equiv. per thiol) was added to the emulsion and the polymerization was allowed 3.5 hours to react. The pH was then lowered to ∼8 with the addition of PBS salts (50 mM final concentration of phosphate) and acetic acid before adding 25 mg of 5-(iodoacetamido)fluorescein (0.049 mmol, 0.025 equiv.s per thiol) in 200 µL of a 1:1 DMF/THF solution followed 10 minutes later by 68 mg of the tetrafunctional cross-linker pentaerythritol tetraacrylate (PA_4_, 0.19 mmol, 0.2 equiv.s to thiol) dissolved in 100 µL of THF. After an additional 1 hour of reaction time, a further 34 mg of PA4 (0.1 mmol, 0.1 equiv.s per thiol) was added in 100 µL of THF; the solution was brought to pH 7.4 with acetic acid and allowed to stir overnight (∼16 hours). The reaction mixture was first filtered through sterile 0.45 µm PES filters and was dialyzed via hollow-fiber dialysis on a Millipore Labscale Tangential Flow Filtration (TFF) System using a Pellicon XL50 Biomax® 100 kDa MWCO membrane until the filtrate had a conductivity equal to that of deionized MilliQ water. The solution was concentrated to approximately 25 mg/mL of PPS-NPs and sterile filtered.

### Donor Organ Procurement

Paired kidneys were procured from n=6 Landrace pigs with a mean dry weight of 80 kg as previously described^9^. All pigs were veterinary inspected and culled in accordance with the European Union Council Regulation (EC) 1099/2009 on the protection of animals at the time of killing. In brief, pigs were rendered unconscious via electrical stunning and sacrificed via exsanguination. Autologous blood was collected into a sterile receptacle containing 100 mL of 0.9% normal saline supplemented with 40,000iU-unfractionated heparin (Fannin, UK). A midline laparotomy was performed, and the kidneys procured en-bloc and immediately placed on ice. The renal artery and ureter were isolated, cannulated and flushed with 20 mL of glyceryl trinitrate (GTN; Hameln, Germany – distributed by Gloucester, UK). Kidneys were randomized to receive a preservation flush with either 1 L 4°C Custodiol (Dr Franz Köhler Chemie Gmbh, Germany). or 1 L 4°C Custodiol supplemented with PPS-NPs to a final concentration 0.25 mg/mL. Both preservation flushes contained 5,000iU-unfractionated heparin and were flushed at a hydrostatic pressure of approximately 100 mmHg. Kidneys were then submerged in either 500 mL Custodiol or 500 mL Custodiol supplemented with 0.25mg/mL PPS-NPs according to their previous randomization. All kidneys were placed on ice for a standardized SCS time of 3 hours.

### Normothermic Machine Perfusion (NMP)

Following 3 hours SCS, kidneys were perfused on separate NMP circuits for 6 hours as previously described ^9^. In brief, the NMP circuit contained a bespoke kidney chamber, a reservoir and extracorporeal membrane oxygenator (Trilly Pediatrics, Eurosets, Italy), a centrifugal pump (ECMOLife, Eurosets, Italy) and an arterial filter (Sherlock, Eurosets, Italy). A heater (Eurosets, Italy) was attached to the oxygenator which was set to a target physiological perfusate temperature. A flow probe, air sensor, temperature probe, pressure line, and an oxygen line (connected to a 95% O_2_ – 5% CO_2_ carbogen gas cannister) were also connected. Prior to connecting the kidneys, the system was primed with approximately 1.4 L perfusate to ensure no air was in the circuit. This consisted of 500 mL Ringer’s solution supplemented with bovine serum albumin (BSA), 800 mL of autologous, cell-saved, leukocyte depleted packed red blood cells (prepared with an Xtra Cell Saver, Livanova, UK) to achieve a target haematocrit of 25-30%. The perfusate was further supplemented with broad spectrum antibiotics (Meropenem), dexamethasone, sodium bicarbonate, glucose, mannitol, and heparin. One circuit was supplemented with antioxidant PPS-NPs (to a final concentration of 0.25mg/mL of NPs). GTN and nutrition were infused into the circuit at a flow rate of 10 mL/hr. A gas mixture of 95% oxygen and 5% carbon dioxide was supplied to the membrane oxygenator. The circulating perfusate temperature was set to 38°C ± 1°C to rapidly re-warm the kidney to a physiological temperature.

Once the circuits were primed, kidneys were removed from ice and flushed with 200 mL of 4°C Ringer’s solution to remove any residual preservation solution. A pre-perfusion sample was taken for reference and kidneys were connected to the circuit via the renal artery. The mean arterial pressure was gradually increased until a target of 75 mmHg was achieved. At this time infusions of GTN and nutrition were commenced. The organ chamber was covered for humidification and to maintain organ temperature. Serial observations and blood gas analyses were recorded and correctable changes to abnormal blood gas physiology were instituted. Ringer’s solution was used to replace the exact volume of urine produced by the kidney.

### Transplant Reperfusion Model with Allogeneic Blood

Bilateral kidneys were retrieved from a further n=6 pigs, as above. Additional blood was also collected per experiment and blood (AO) matched to the donor pig using DiaClon Anti-A (Bio-Rad, Germany). Kidneys were perfused via NMP as above before being transferred to new NMP circuits that were primed with the AO matched blood to represent the ‘recipient’.

#### NMP Transplant Model

Following preservation with NMP, kidneys were removed and flushed with preservation solution before being placed back on ice for 1 hour. During this time two new NMP circuits were primed with 1.2 L whole blood from a blood matched, genetically different donor pig to mimic the transplant procedure upon reperfusion, without immunosuppression. The only supplements were 250 mg Meropenem and 10 mL 15% glucose. No PPS-NPs were included in this part of the study. A syringe driver infused nutritional support (10 mL/hour). After 1-hour on ice, kidneys were flushed with 200 mL of 4°C Ringer’s solution before being re-weighed and attached to the matched-blood NMP circuits. Both kidneys were perfused for 6 hours at a target mean arterial pressure of 80 mmHg.

### Sample Collections

#### Perfusate

Serial perfusate samples were collected. All samples were centrifuged at 1000 g for 10 minutes at 4°C. The supernatant was decanted into 1 mL aliquots and stored at -80°C.

#### Urine

Serial urine samples were collected and aliquoted into 1 mL cryovials and stored at -80°C.

#### Biopsies

At the end of perfusion, the weight of kidneys was recorded, and wedge biopsies were stored in 10% neutral buffered formalin (Merck, Germany).

#### Infra-Red (IR) Imaging

IR images were taken of the kidneys to determine perfusion homogeneity/heterogeneity. Images were taken prior to the kidneys being connected to the circuit immediately following cold storage, followed by 5-minute intervals for the first 30 minutes. Serial images were then taken until the end of the experiment. The entire surface of the kidney was populated with temperature spots and the mean and standard deviation temperature was used as a measure of perfusion heterogeneity. For each time-point the emissivity was set at 0.98, reflective temperature at 22°C, distance 1m and the relative humidity set at 50.0%.

#### Biochemical Analysis

A blood gas analyzer (GEM 4000, Werfen, UK) was used to assess pH, pO_2_, pCO_2_, electrolyte concentrations and Co-Oximetry within the circuit, which were corrected to maintain physiological levels.

### Biochemical Assays

#### Urine total protein and albumin content

Urine samples (80 µL) were centrifuged at 800 g for 10 minutes at 0°C and the supernatant diluted 1:20 with MilliQ water. Total protein content was quantified using Pierce^TM^ BCA protein assay kit as per the supplier’s protocol (Thermo Scientific, USA). For albumin, the supernatant was diluted 1:25000 or 1:50000 and quantified using an Albumin pig ELISA kit (Abcam, Cambridge, UK) according to manufacturer’s instructions.

#### Oxidized DNA

Quantification of oxidized DNA was performed using a DNA Damage Competitive ELISA kit (Invitrogen, USA) according to manufacturer’s instructions with plasma samples diluted 1:8 in assay buffer.

#### TNF-α

Quantification of TNF-α was performed on undiluted samples using a porcine TNF-α ELISA kit (Invitrogen, USA) with 20 pg/mL sensitivity according to manufacturer’s instructions. Samples that were determined to be below the detection limit were quantified using <0.3 pg/mL sensitivity porcine TNF-α ELISA kit (Invitrogen, USA).

### Complement (sC5b9) Assay

Soluble C5b-9 was quantified by MesoScale Discovery (MSD). MSD GOLD 96-well streptavidin SECTOR plate was coated with 2 μg/mL biotinylated anti-C5b-9 antibody (Abcam) and blocked with PBS containing 2% BSA 0.05% Tween 20 (v/v). Kidney perfusate was diluted three-fold and added to the plate. Human sC5b-9 (Complement Technology) at concentration ranging from 5 μg/mL to 0.01 μg/mL was used to generate a standard curve. Kidney samples and sC5b-9 standard were diluted in PBS containing 2% BSA, 0.05% Tween 20 and 10 mM EDTA. Ruthenylated anti-C6 monoclonal antibody (2 μg/mL) diluted in MSD Diluent 100 was used for detection. Plate was read using an MSD Sector Imager 6000 and sC5b-9 concentration was quantified from interpretation of electrochemiluminescent (ECL) signals using MSD Discovery Workbench 4.0.12.1 software.

### Histological Analysis

#### Haematoxylin and Eosin

Kidney tissue biopsies that were fixed in 10% buffered formalin solution were dehydrated and paraffin-embedded. Sections were cut (5 mm), de-paraffinized and stained with hematoxylin and eosin using the protocol set by the Leica Autostainer XL (Leica Biosystems Nussloch GmbH, Nussloch, Germany). Samples were graded using the Remuzzi scoring system by a clinical histopathologist who was blinded to the sample groups. Histologic morphological changes were observed in all kidneys at the end of the 6-hour reperfusion, although these were observed to be only acute features. Overall, there was better preservation of tissue architecture in PPS-NP-treated kidneys, with less overall acute tubular necrosis (Supplementary Table 1). Histology was statistically analyzed using a one sample Wilcoxon t tests with significance displayed with respect to healthy kidney score (0).

#### Immunostaining

Kidney sections were incubated in 30% sucrose solution in 0.005 M PBS for 72 hr prior to being sectioned into 40µm slices using a freezing cryostat. The sections were permeabilized with a 0.3% v/v Triton X-100 in PBS (PBS-T) for 30 minutes at room temperature followed by a blocking solution (2% normal goat serum in PBS-T) for 1 hour at room temperature. Tissue sections were washed three times with PBS-T, incubated with VCAM-1 monoclonal antibody (1:100, 1.G11B1 Invitrogen) at 4°C overnight, followed by incubation with goat anti-mouse Alexa 546 (1:500; Invitrogen) for 2 hours at room temperature and counterstained with Hoechst (1:1000; Sigma-Aldrich) for 10 minutes prior to imaging. The stained tissues sections were mounted using ProLong®Antifade mounting reagent (Thermo-Fisher Scientific) and imaged using a Leica SP5 confocal microscope (Wetzlar, Germany) with 40X magnification.

### Statistical Analysis

Statistical analysis was performed using Graphpad Prism. Data normality was formally assessed using the Shapiro-Wilk test. Unless otherwise stated, a two-way ANOVA (mixed-effect analysis) with the Fishers Least Significant Difference (LSD) test for multiple comparisons was used to assess significance between groups and individual timepoints respectively. Data was analyzed pairwise between untreated and PPS-NP treated kidneys from the same pig. Error bars represent +/-SEM.

For the complement assays a mixed model was fitted to the log transformed sC5b-9 concentration for each time point. The model terms were: *response variable*: log_10_ concentration, *fixed effect model*: kidney, *random-effect model*: animal/kidney/replicate.

The model term ‘kidney’ also represents the treatment effect, as the two treatments (with and without PPS-NP) were applied to two paired kidneys. A significance test was performed as part of the model-fitting process to determine whether the mean with/without PPS-NP ratio was different from 1 (or equivalently, mean log_10_ ratio concentration was different from 0).

## Results and Discussion

DGF represents a major clinical challenge following transplantation and is associated with poor long-term graft function and patient survival (estimated to be a 40% decrease in long-term survival).^30^ IRI significantly contributes to DGF, with prolonged ischemic times leading to an exacerbation of acute kidney injury and an increased risk of primary non-function.^31^ There have been attempts to reduce IRI which include improvements to donor and recipient management, as well as therapeutic interventions (for example, calcium-channel blockers, L-arginine and N-acetylcysteine), but with limited clinical success.^32^ Indeed, there are still no specific therapies available in the clinic to reduce IRI following transplantation. A primary challenge in managing IRI is that graft ischemia is ongoing during preservation. Therefore, treating the donor kidney prior to transplantation may mitigate intra-operative and post-transplant induced injury. With the advent of NMP, there is an opportunity to directly treat the donor kidney prior to transplantation, specifically targeting pathways associated with IRI, with reduced off-target or systemic risks to the recipient. In this study, we have therefore evaluated if 1) direct treatment of a donor kidney with antioxidant PPS-NPs could be used during preservation and NMP to improve perfusion parameters and 2) determine if this ameliorates post-transplantation IRI (**Scheme 3**).

**Scheme 3.**
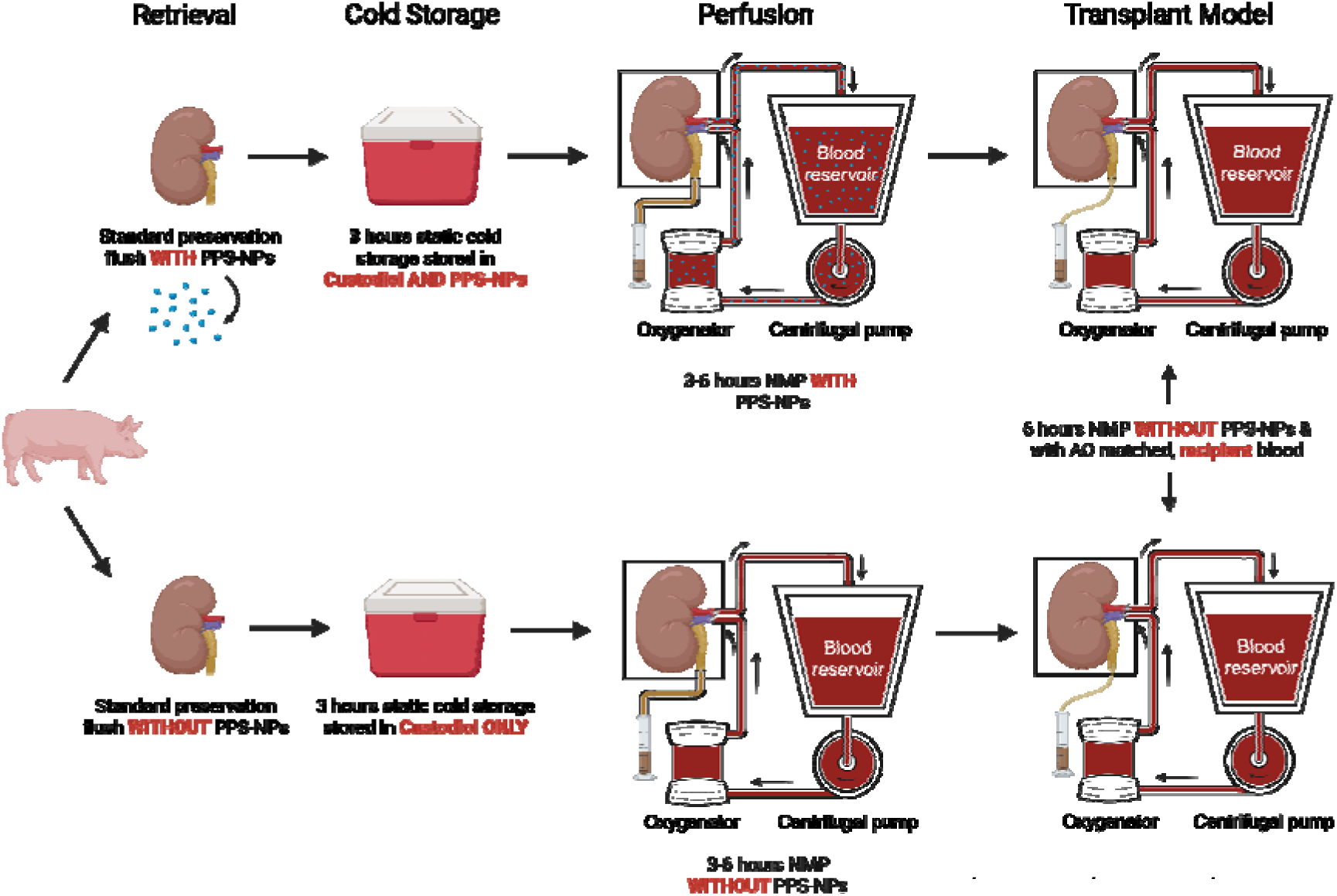
Procedural setup of kidney harvesting and storage and simplified experimental design of NMP and transplant perfusion circuits. Paired kidneys were retrieved from the same donor with one treated with PPS-NPs and the other acting as the matched untreated control.

### Perfusion (NMP)

We successfully synthesized 6.5 g of crosslinked PPS-NPs with a Z-Ave size of ∼150 nm with a narrow PDI of 0.091 (**Figure S1**). Previous studies have demonstrated their exceedingly low toxicity and anti-inflammatory action.^33–36^ All kidneys were successfully perfused for 6 hours, achieving average renal blood flows (RBF) of 84±38.92 mL/min/100g in control kidneys and 102±31.86 mL/min/100g in PPS-NP treated kidneys. Hemodynamic measurements confirmed that NMP was successfully able to perfuse kidneys from both groups over a 6-hour period at a constant mean arterial pressure of ∼75 mmHg (**Figure 1A**). Interestingly, kidneys that were stored via SCS and then exposed to NMP with PPS-NP supplementation (final solution at 0.25 mg/mL PPS-NP) had an overall 17% higher average RBF (61.38±17.03 vs 71.86±9.83 mL/min/100 g, **Figure 1B**). This is likely explained by the significantly lower intra-renal resistance (IRR) found in the PPS-NPs supplemented group, particularly at immediate, and later time points, predominantly within the 3–6-hour range (**Figure 1C**). Together, these data indicate PPS-NPs are facilitating a more homogenous flow of blood through the kidney. Blood vessels with a low flow due to higher resistance may also experience more damage or inflammation. Indeed, it is well documented that patients that experience poor renal haemodynamics in the early post-transplant period are likely to develop DGF which impacts graft survival.^37, 38^

**Figure 1.**
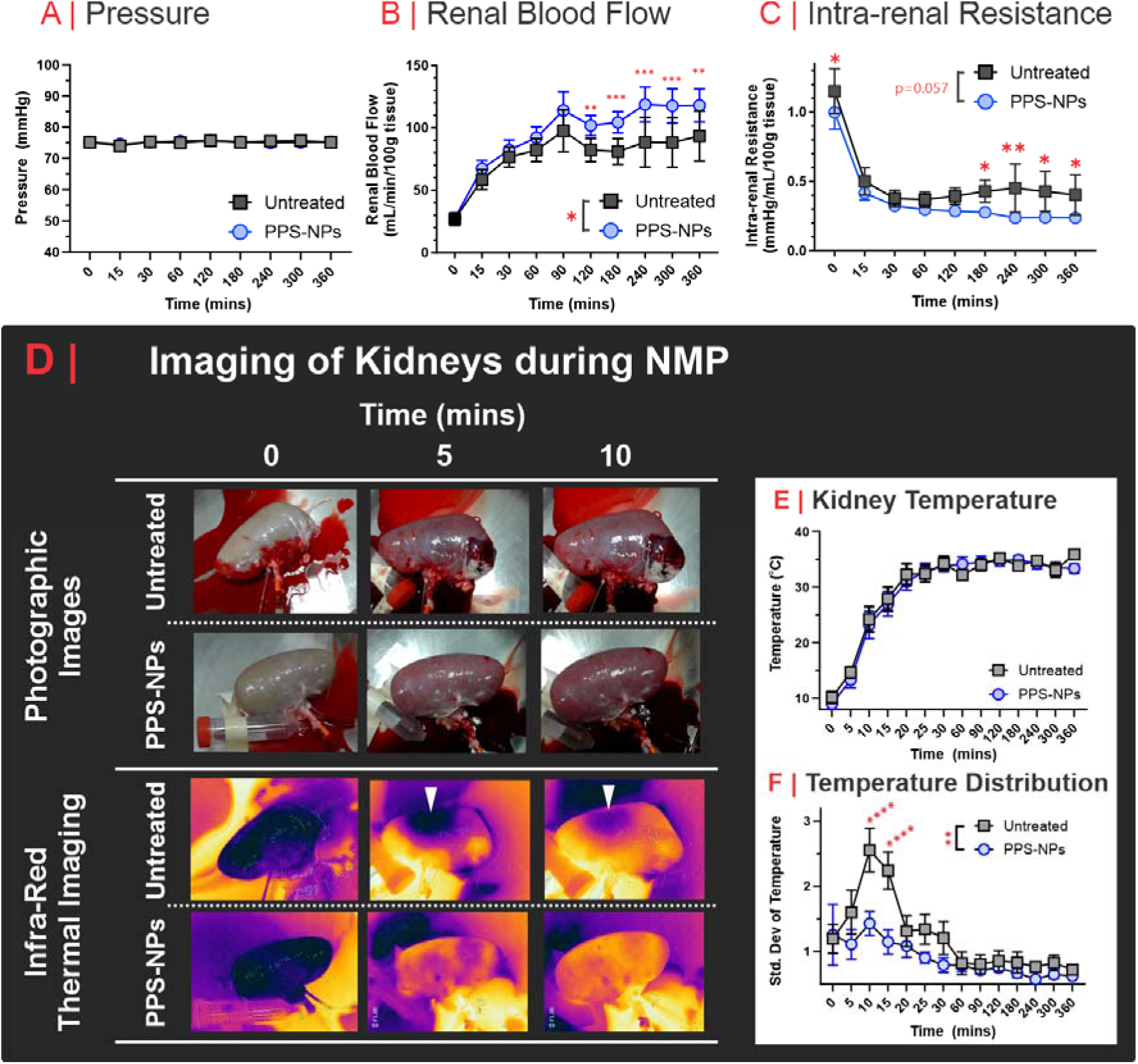
Top: Hemodynamical analysis of the kidney was determined by measuring **A**. renal pressure, **B.** renal blood flow and **C.** intra-renal resistance for both PPS-NP supplemented and non-supplement NMP using leukocyte depleted autologous blood; *; P≤0.05,**; P≤0.01, ***; P≤0.001. Lower: temperature analysis of kidney during NMP of cold stored kidney. Representative **D.** digital photographs and infra-red thermal imaging of the kidney during the first 10 minutes of NMP. White triangles indicate heterogenous warming of the kidney (as cold patches) within the untreated perfused kidneys and is indicative of poor blood flow. Infrared imaging yielded **E.** average kidney temperature and **F.** temperature distribution over time. Data represented as an average of 6 kidneys per group. **; P≤0.01, ****; P≤0.0001.

Whilst haemodynamics were significantly improved with PPS-NP, there was no difference in the average rate of rewarming (**Figure 1D-E**); however, IR analysis indicated that PPS-NPs resulted in a significantly more homogenous and reproducible warming of the kidney (**Figure 1F**, p=0.002), with this most evident between 5 and 15 mins. In the IR images, cold spots were evident in the untreated kidneys (white arrows, **Figure 1D**) potentially indicating low blood flow in these localized areas. Kidneys supplemented with PPS-NPs did not display such cold spots, indicating improved haemodynamics and vascular tone with respect to unsupplemented NMP.

In both groups, kidneys demonstrated excellent control of biochemical parameters and maintained homeostatic control of pH (**Figure 2A**) with little need for additional sodium bicarbonate to manage acidosis (**Figure 2B-C**). Furthermore, bicarbonate, calcium and potassium all remained within or close to their physiological ranges for both experimental groups (no statistical differences, **Figure 2**). Sodium gradually increased during NMP, likely a result of Ringer’s supplementation to replace urine output. Likewise, glucose increased over time, however this is likely due to renal gluconeogenesis (**Figure 2D**). Interestingly, lactate was ∼2x lower in the perfusate of PPS-NP treated kidneys compared to control, between 240-360 minutes (**Figure 2E**). Given that lactate is a surrogate marker of anaerobic metabolism and tissue perfusion, this further indicates hypoperfusion in control kidneys while further supporting the use of antioxidants to improve vascular integrity and reduce IRI related inflammation. No differences were observed between pre-and post-perfusion kidney weight, with untreated kidneys increasing on average by 6±12.5% and PPS-NP treated kidneys by -2.5±7.9%.

**Figure 2.**
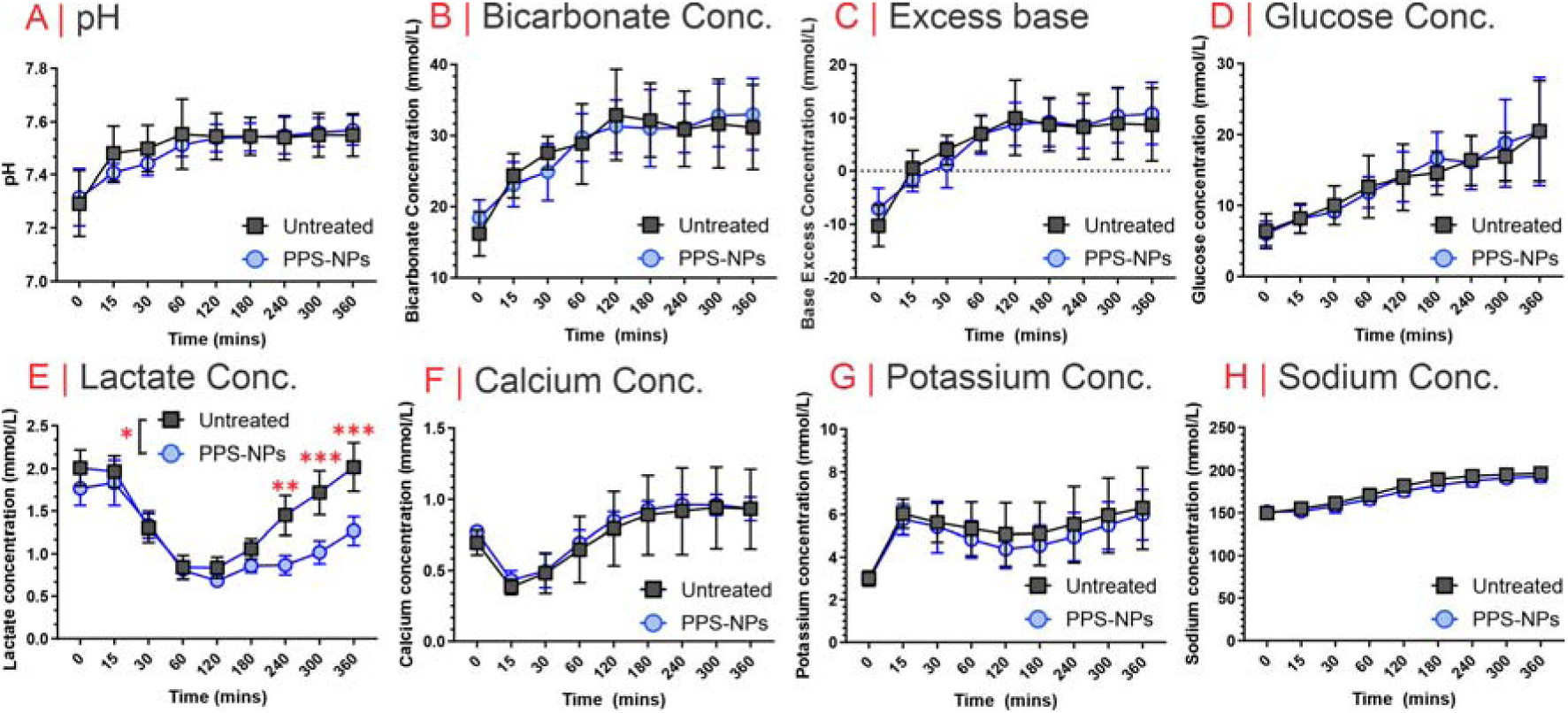
Biochemical analysis of kidney perfusates including **A.** pH analysis, **B.** bicarbonate concentration, **C.** excess base, as well as the concentration of **D.** glucose, **E.** lactate, **F.** potassium, and **H.** sodium. Statistical significance between groups was determined using a Two-Way ANOVA (mixed-effect analysis) with the Fishers Least Significant Difference (LSD) test for multiple comparisons of individual points; *; P≤0.05,**; P≤0.01, ***; P≤0.001.

One of the key parameters for donor kidney functionality is urine output. As such, the volume of urine and its contents were assessed to evaluate filtration function (**Figure 3**). We found that both kidneys almost instantly began to produce urine, with the control kidneys producing significantly more urine (p<0.05) than the PPS-NP supplemented (**Figure 3A**). Interestingly, total protein content and albumin concentrations in urine, biomedical indicators of improper renal function, were both significantly higher in the control kidneys, with values up to ∼75% higher than those of PPS-NP treated kidneys (**Figure 3B-C**). This suggests that although the control kidneys produced more urine, this may be due to either damage, inflammation, or stress in the kidney permitting the atypical passage of large molecular weight proteins such as albumin into the urine (proteinuria). Note, ordinarily albumin is too large to pass through renal filtration structures and is actively removed from the urine. Elevated levels, particularly in the control kidney urine could potentially be explained by more heterogenous blood flow through the kidney which may create areas of localized high pressure, permitting passage of molecules above the renal exclusion limit into the urine.

**Figure 3.**
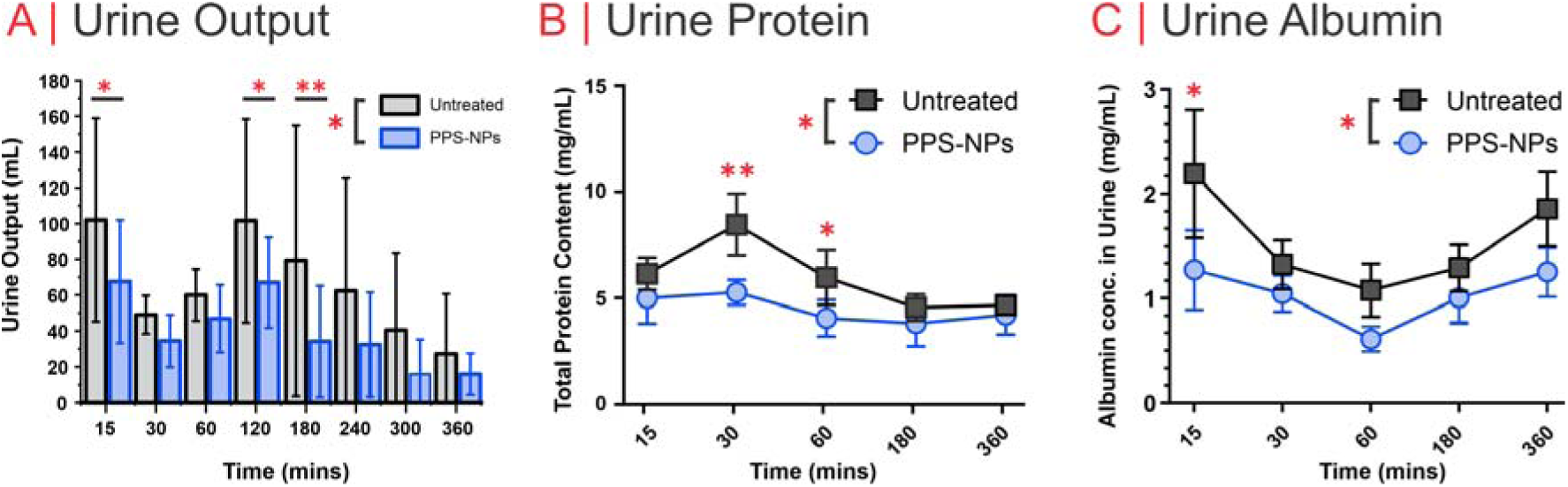
Urinalysis was performed to assess kidney function with **A.** urine output volume**, B.** urine total protein, and **C.** urine albumin measured. *; P≤0.05,**; P≤0.01, ***; P≤0.001.

### Transplant Model

It is well documented that the condition of the donor organ prior to transplantation directly correlates with clinical outcomes.^39^ To evaluate the post-transplant impact of PPS-NP treatment, we utilized a transplant reperfusion model. New perfusion circuits were primed with whole blood from AO-matched allogeneic donors without any PPS-NP supplementation. From precluding perfusion (NMP) experiments above, minimal differences in hemodynamic, biochemical, and urine analysis readouts between 3 and 6 hours (**Figures 1-3**) were observed, with most parameters stabilizing by 3 hours. Additionally, untreated kidneys began to show signs of stress after 3 hours, as evidenced by increasing lactate concentrations (**Figure 2**). Therefore, for subsequent experiments, we transitioned to the transplant circuit after 3 hours on the perfusion (NMP) circuit. By minimizing damage in the untreated kidneys, we thereby set a higher benchmark for the PPS-NP treatment to demonstrate its efficacy in the next transplantation phase. Nevertheless, 3 hours is ample time for ’resuscitating’ cold-stored donor kidneys on the NMP circuit and ensuring any IRI-associated damage occurs in this highly mitigated setup.

As with NMP, haemodynamics of PPS-NP treated kidneys were significantly improved compared to the untreated after allogeneic reperfusion in this transplant circuit. At a constant pressure of ∼80 mmHg, the PPS-NPs had an RBF that was up to 2x higher and an IRR that was on average 2x lower than that of the untreated control kidneys (**Figure 4A-C**), indicative of improved vascular tone and endothelial integrity. This is likely explained by the higher inflammatory activation within the untreated kidneys restricting the blood flow within the blood vessels. Indeed, histological (H&E) sections of PPS-NP treated kidneys presented with better tissue integrity and reduced acute tubular necrosis as adjudicated by a treatment blinded histopathologist (**Figure 4D** and **Table. S1**). Damage to the donor kidney prior to transplantation is causally linked to DGF and a reduction in 1 year survival rates.^40^ As such, improving tissue integrity during preservation is of significant clinical importance.

**Figure 4.**
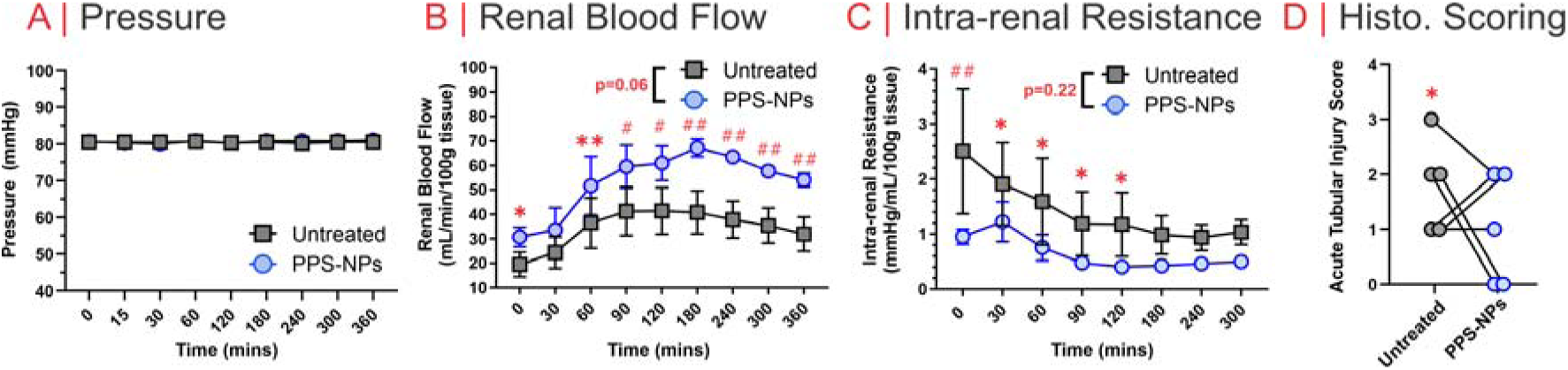
Transplant reperfusion model haemodynamics in allogeneic blood. **A.** blood pressure, **B.** renal blood flow, and **C.** intra-renal resistance. **D.** Histological scoring of paired kidneys (from the same pig) by a study blinded histopathologist; *; P≤0.05,**; 0.001<P≤0.01,^#^; 0.0001<P≤0.001,^##^; P<0.0001.

Next, we assessed the kidney’s ability to filter blood and maintain homeostasis within healthy physiological ranges. Here, we found that sodium bicarbonate, base and pH all increased at equal rates in both groups until the end of reperfusion while predominantly remaining within physiologically healthy ranges reported for pigs (**Figure 5A-C**). The regulation of critical electrolytes in the blood such as calcium and sodium remained stable and physiological in both groups, whereas glucose and lactate were elevated but declined during allogeneic reperfusion, falling within the physiological range over the course of this experiment (**Figure 5D-G**). There was a marginal decline in lactate concentration over time from an average of 5.5±3.0 to 4.7±2.6 mmol/L in untreated kidneys and 5.4±2.9 to 3.5±1.2 mmol/L in PPS-NP-treated kidneys, but both remained within physiological ranges from 30 minutes onwards (**Figure 5E**). The decline in lactate in both groups demonstrates active biological processing via the Cori Cycle. Urine output was significantly higher in PPS-NP treated kidneys at 30 minutes, but urine production in both kidneys diminished over the next 6 hours (**Figure 5H**).

**Figure 5.**
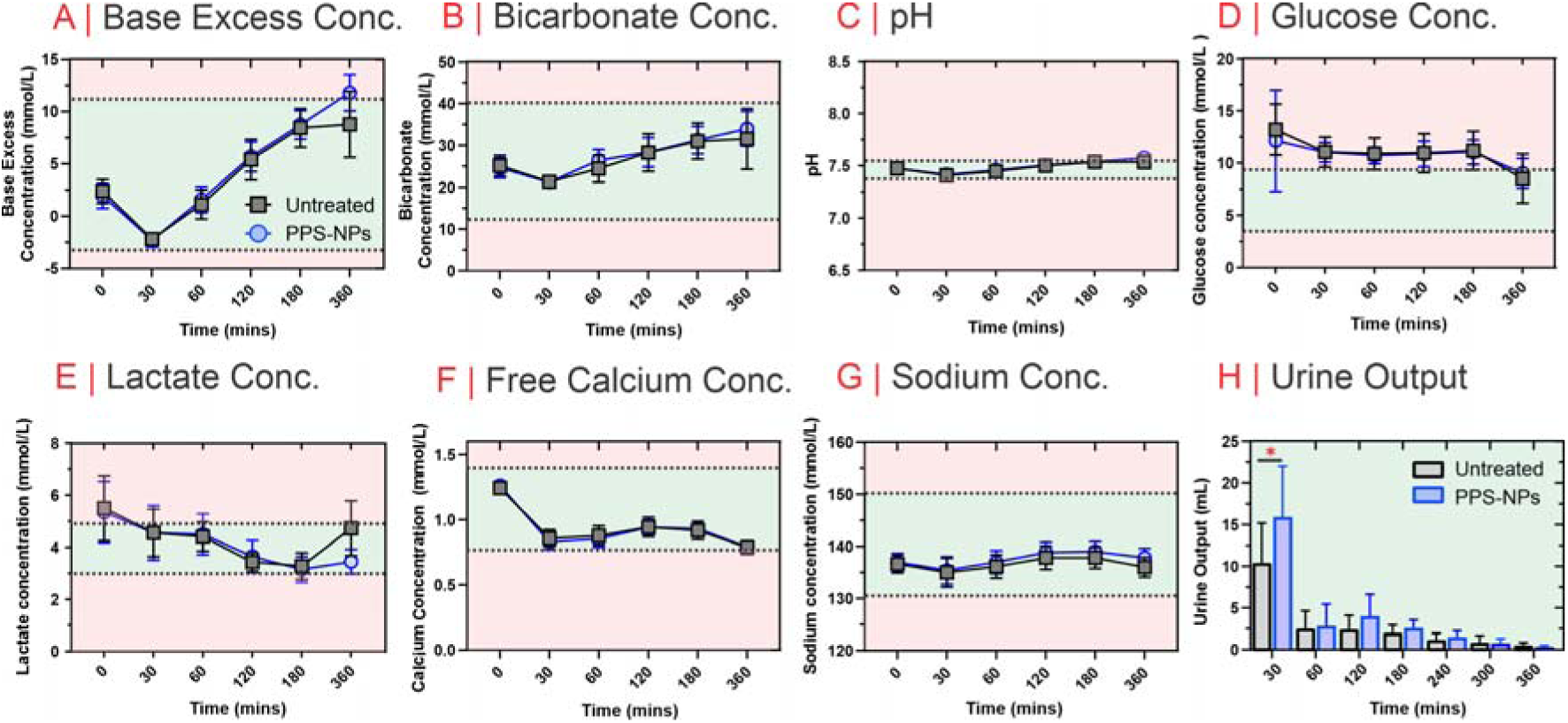
Kidney function parameters during the transplant reperfusion model with allogeneic blood. Biochemical readouts include **A.** base excess, **B.** bicarbonate, **C.** pH, **E.** lactate, and **F.** free (iodized) calcium concentration, and **H.** urine output. In **A-G**, physiological range for healthy pigs is highlighted in green (outside range in pale red).

The remarkable improvements in haemodynamics in both the NMP and transplant circuits observed in this study underscore the potential of PPS-NPs in mitigating DGF and enhancing transplant outcomes. The hemodynamic benefits noted, including enhanced RBF and decreased IRR, suggest a complex interplay of underlying mechanisms leading to blood vessel inflammation and occlusion. This directs our focus toward the molecular interactions influenced by PPS-NPs, particularly oxidative stress/tissue damage, endothelial inflammation, and complement activation, and how they impact upon renal tissue health during NMP and transplantation.

### Mechanistic Insights: PPS-NPs are anti-inflammatory and block complement activation

For this study, the PPS-NPs were selected as the antioxidant based on their potent ROS-scavenging capability (particularly for H_2_O_2_, hypochlorite, hydroxyl radical and peroxynitrite) as well as their greater stability than biological therapeutics (e.g., catalase). Their scavenging action is hypothesized to confer significant therapeutic benefits in the context of IRI associated with transplantation. The rationale behind this lies in the detrimental roles that ROS plays in transplant organs where it collectively contributes to the oxidative damage of critical biomolecules, including proteins, lipids, and nucleic acids (DNA), and are implicated in triggering inflammatory responses, including complement activation.^41, 42^ Injury to transplant grafts during ischemia, particularly at the point of reperfusion, is predominantly driven by aberrant formation of ROS.^43^ It is well-established that an imbalance characterized by excessive ROS production and diminished antioxidant defenses precipitates cell membrane damage, apoptosis, necrosis, and, consequently, DGF or even failure.^44^ This study posits that PPS-NPs may confer protective effects against IRI through mechanisms of 1) direct ROS scavenging and 2) inhibition of the complement system, both of which play critical roles is graft failure.

An evaluation was therefore conducted to assess the efficacy of antioxidant PPS-NPs, in mitigating oxidative damage to transplanted kidneys. To quantify the extent of oxidative damage/oxidative stress, we first measured levels of oxidized DNA (8-Oxo-dG), during NMP. This phase is critical as it involves warming and reoxygenation of the kidney post-cold storage, where the main insult of IRI it thought to occur. It was determined here that the PPS-NPs were able to significantly reduce the level of oxidized DNA with respect to the untreated control kidneys (p≤0.05), with a 5.3x lower concentration after 3 hrs of autologous perfusion (p≤0.001) (**Figure 6A**). These data appear to confirm our hypothesis that PPS-NPs are able to scavenge ROS during cold storage and NMP, alleviating much of the oxidative stress associated with NMP. This is further evidenced by the downstream inflammatory cytokine, TNF-α, which also undergoes a significant (1.6x, p≤0.05) reduction in concentration after 3 hrs in the PPS-NP treated group (**Figure 6B**). These data corroborate previous works that have demonstrated PPS-NPs reduce TNF-α levels in various inflammatory disease models.^26, 34, 45^

**Figure 6.**
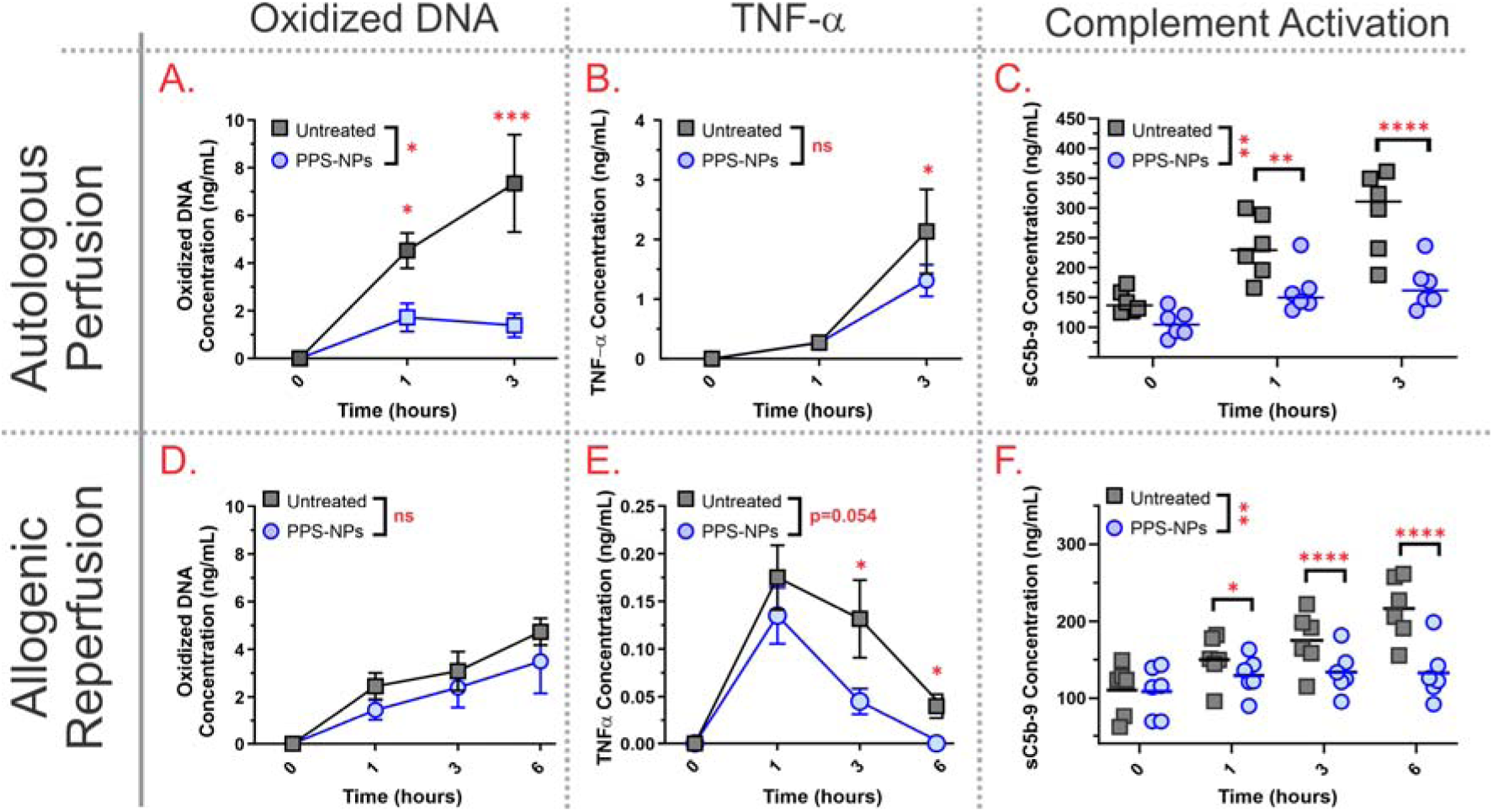
Kidney damage, oxidative stress, and inflammatory markers in both NMP (autologous blood perfusion) and during the transplant reperfusion model (allogeneic blood reperfusion). Blood concentrations for **A.**/**D.** oxidized DNA (8-Oxo-dG), **B.**/**E.** TNF-α, and **C.**/**F.** complement component sC5b-9 were determined via ELISA for autologous perfusion/allogeneic reperfusion respectively. *; P≤0.05,**; P≤0.01, ***; P≤0.001, ****; P≤0.0001.

Although not completely understood, aberrant complement activation also ensues after transplantation, with the formation of the terminal cytolytic membrane attack complex (C5-9b, also referred to as the membrane attack complex, MAC), inducing direct renal injury via apoptosis of epithelial tubular cells^46^. During NMP, there was a significant reduction in soluble MAC (sC5-9b) in the perfusate of PPS-NP treated kidneys (p≤0.01), with a 1.72x lower concentration of sC5-9b in the PPS-NP treated kidneys at 3 hrs (p≤0.0001) (**Figure 6C**). This result is likely rationalized by the fact that complement can be both directly and indirectly activated by ROS.^47, 48^ Direct activation occurs through the oxidation of reactive thioester bonds between C3b and C5b, releasing C3a and C5a respectively. Indeed, it has also been found that antioxidants such as catalase^47^ and methionine^48^ (for which PPS-NPs are biomimetic of) were able to inhibit this direct activation pathway. Oxidative stress also indirectly increases complement activation through both an upregulation of inflammatory pathways, but also through chemical (oxidative) modification of glycoproteins on cell surfaces^49, 50^, lipids^51, 52^ or proteins more broadly^53^, that can render them complement activating through either classical (due to neoantigen/neoepitope formation) or alternative activation pathways (due to increased nucleophilicity). Additionally, endothelial cells experiencing oxidative stresses after ischemia have been found to have aberrant surface presentation of cytokeratins which are potent binders of mannose-binding lectin (MBL) and thus strongly activate the lectin pathway of complement activation.^54^ Notably, cytokeratin autoantibodies have also been found in the blood of transplant patients, corroborating this pathway of complement activation in a clinical transplant setting.^55^ These highlight the potential of antioxidant therapies to work across multiple inflammatory pathways and severely limit graft damage and rejection.

When looking at the transplant model, we only see a mild increase in oxidized DNA, with the levels in the PPS-NP treated kidneys always lower, albeit not statistically significant (**Figure 6D**). The absence of PPS-NPs in the transplant circuit likely explains the lack of significant differences observed here. Additionally, very low TNF-α levels, even in the untreated controls, underscore the effectiveness of NMP in mitigating severe kidney damage during IRI (**Figure 6E**). Interestingly, despite the presence of allogeneic leukocytes in the reperfusion blood, TNF-α concentrations were significantly lower - an order of magnitude less than those seen in NMP with autologous leukocyte-depleted blood. Nevertheless, TNF-α levels in PPS-NP treated kidneys were approximately three times lower at 3 hours than untreated controls (p≤0.05) and had undetectable levels at 6 hours (**Figure 6E**). This suggests a markedly reduced inflammatory profile in PPS-NP treated kidneys, which could point to a decreased incidence of DGF and improved transplant outcomes. However, it should be noted that control kidneys also showed extremely low and decreasing TNF-α levels, further demonstrating NMP’s effectiveness in enhancing transplant quality and success rates.

Similarly to the NMP circuit, complement activation (sC5b9) levels in the untreated blood-perfusate from the transplant reperfusion circuit approximately doubled from baseline over 6 hours in allogeneic blood. Remarkably, sC59b levels in the PPS-NP treated kidneys remained at baseline levels for the entire 6-hour transplant experiment (**Figure 6F**).

To interrogate more directly the inflammatory state of the blood vessel, we performed immunohistochemistry on the glomerulus using VCAM-1 as a surrogate marker of inflammatory activation and blood vessel damage. Mechanistically, VCAM-1 is important for the ‘firm’ binding of circulating leukocytes and high levels would be indicative of strong leukocyte attachment (**Scheme 1**).^56, 57^ In this regard, neutrophil rolling is known to occur on the vascular endothelium, along with vascular wall remodeling, ultimately leading to narrowing of the lumen and reduced passage of blood flow, and DGF (**Scheme 1**).^58^ Minimizing blood vessel inflammation would therefore be advantageous in reducing the risk of an immune response against the donor kidney and ultimately lowering the risk of transplant failure. In the NMP circuit, we observe higher IHC staining levels of VCAM-1 in the untreated controls with respect to the PPS-NP treated samples, clearly outlining arterioles of the glomerulus (**Figure 7**). The more intense VCAM-1 in the control kidneys correlated with higher levels of plasma inflammatory and damage markers such as oxidized DNA, TNF-α, and sC5b9. Critically, this would suggest that unlike soluble markers which are flushed before transplantation, a fixed inflammatory signature of the kidney blood vessels, such as VCAM-1, will be carried over into the transplant recipient, increasing the risk of transplant failure.

**Figure 7.**
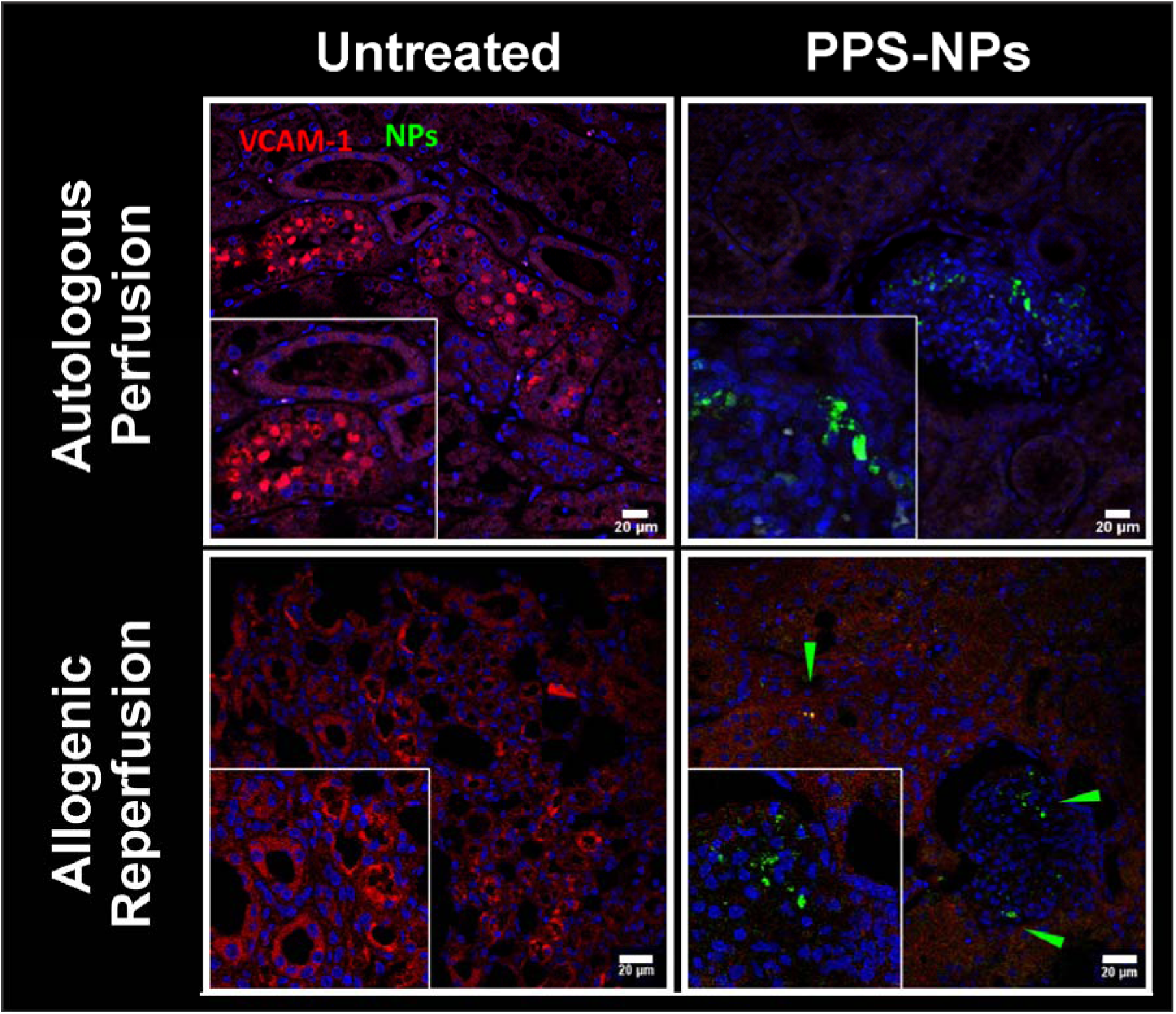
Immunohistochemistry of kidney glomerulus biopsies in (Top) NMP autologous blood perfused and (Bottom) transplant reperfusion model, either (Left) untreated or (Right) PPS-NP treated. Nuclei are stained with Hoechst (blue), inflamed arterioles are stained using a VCAM-1 antibody (red), and PPS-NPs are fluorescently labelled with 5-(iodoacetamido)fluorescein (green). Green arrows indicate the location of a PPS-NP cluster.

Similarly to NMP, the presentation of leukocyte adhesions receptor, VCAM-1, was lower in the PPS-NP treated kidneys (**Figure 7**). Interestingly, despite most PPS-NPs being flushed from the kidneys, small clusters could be found sparsely located (green arrows), with VCAM-1 staining appearing to negatively correlate with the PPS-NP clusters. This observed reduction in VCAM-1 with PPS-NPs treatment across both the NMP and transplantation circuit, demonstrates their ancillary role in reducing leukocyte adhesion molecule presentation. A potential explanation may be that PPS-NPs’ inherent ability to reduce oxidative stress, MAC formation, and TNF-α production, which are all upstream activators of adhesion molecule expression and presentation, reduces VCAM-1, neutrophil rolling, and blood vessel narrowing. Crucially, by protecting the graft/endothelial cells from oxidative damage during cold storage and NMP, this decreases the inflammatory propensity and inflammatory carryover into the transplant.

Of note, TNF-α has been a therapeutic target in the transplant setting for decades, due to its pleiotropic roles in rejection, but to date, there have been no successful reports of anti-TNF-α treatment.^59, 60^ This highlights that for a successful IRI therapy, functionality across multiple inflammatory pathways is paramount. In this study, we demonstrate that antioxidant PPS-NPs do indeed work across multiple levels, including upstream where it is able to disrupt much of the initiating oxidative stimulus, while also dampening or diminishing the inflammatory environment.

In summary, PPS-NPs significantly increased renal blood flow and decreased intrarenal resistance (**Figure 4**), both clinically meaningful indicators of transplant success. Subsequent mechanistic studies demonstrated that reduction in MAC, oxidized DNA, and TNF-α (**Figure 6**) likely led to these enhancements in renal haemodynamics. We hypothesize that the higher blood vessel inflammation (VCAM-1) and complement activation (sC5-9b) in untreated controls results in blood vessel occlusion via increased neutrophil rolling (**Scheme 1**). A further explanation for this is maybe through increased fibrin(ogen) deposition. Other NMP studies have demonstrated that fibrin(ogen) deposition occurs in transplant organs leading to blood vessel obstruction, with fibrinolytic factors found to significantly improve blood flow.^14^ Given complement’s ability to orchestrate fibrin(ogen) deposition, we hypothesize that increased complement activation and MAC formation may also contribute to blood vessel occlusion through this mechanism.^61, 62^ Another potential source could also be from neutrophil extracellular traps (NETs)^63, 64^; minimizing neutrophil rolling, extravasation, and activation would also significantly reduce this possibility. Thus, PPS-NP’s ability to block both complement activation/deposition and leukocyte adhesion molecule presentation makes them an exciting treatment modality with much potential for boosting transplant success rates and rejuvenating marginal organs to a transplantable status. With the critical shortage of donor organs deemed a public health crisis, increasing the number of usable donor kidneys has become a matter of urgency worldwide. This combination of NMP and PPS-NPs thus represents a promising approach to combat this shortfall as it not only mitigates the IRI, oxidative stress, and inflammatory activation, which that often lead to transplant failure, but it also allows an effective real-time assessment of organ suitability before transplantation through indicators such as renal blood flow, intra renal resistance and blood biochemistry.

## Conclusions

PPS-NP therapy appears safe during SCS, NMP, and during transplant reperfusion with allogeneic blood. PPS-NP treatment was associated with improved tissue perfusion and haemodynamics, potentially via suppression of complement activation, oxidative stress, and inflammatory pathways. This was evidenced by a reduction in the soluble form of the terminal cytolytic MAC, (sC5-9b), circulating TNF-α, and leukocyte adhesion molecule, VCAM-1. Given that DGF is common post-transplant, further work evaluating nanoantioxidant therapies as a treatment strategy for IRI is warranted.

By potentially expanding the pool of viable organs, this approach could significantly improve transplantation success rates and patient outcomes. This research sets the stage for future investigations into the optimization of nanoparticle formulations and their application in NMP protocols. The broader implications of this work could extend to the transplantation of other organs, promising a new horizon in transplant medicine. It is our hope that these findings will encourage a re-evaluation of current organ preservation strategies and catalyze the development of more advanced, nanoparticle-augmented perfusion techniques.

## Funding

This work was part funded by a Prize Post-Doctoral Fellowship in collaboration with GlaxoSmithKline (GSK).

## Disclosure

The authors declare no conflicts of interest

## Author Contributions

JPS – Conceptualization, methodology, formal analysis, investigation, writing (original draft), and visualization.

RD – Methodology, formal analysis, investigation, resources, and writing (review and editing).

AG – Methodology and Investigation

KA – Methodology and Investigation

AMF – Investigation and Resources

DD – Resources and data curation

MG – Resources

NG – Investigation, resources, and data curation

GC – Data curation

JB – Investigation

NG – Data curation

MR – Investigation, supervision, and funding acquisition

NT – Investigation, resources, data curation and writing (review and editing)

JEF – Conceptualization, methodology, formal analysis, investigation, writing (original draft), supervision and funding acquisition.

## Notes

### Competing Interest Statement

The authors have declared no competing interest.

### Summary of Updates

The manuscript has been reformatted to include the results and discussion in one main section.

