## Supplementary information for "Antioxidant Polysulfide Nanoparticles Ameliorate Ischemia-Reperfusion Injury and Improve Porcine Kidney Function Post-Transplantation"

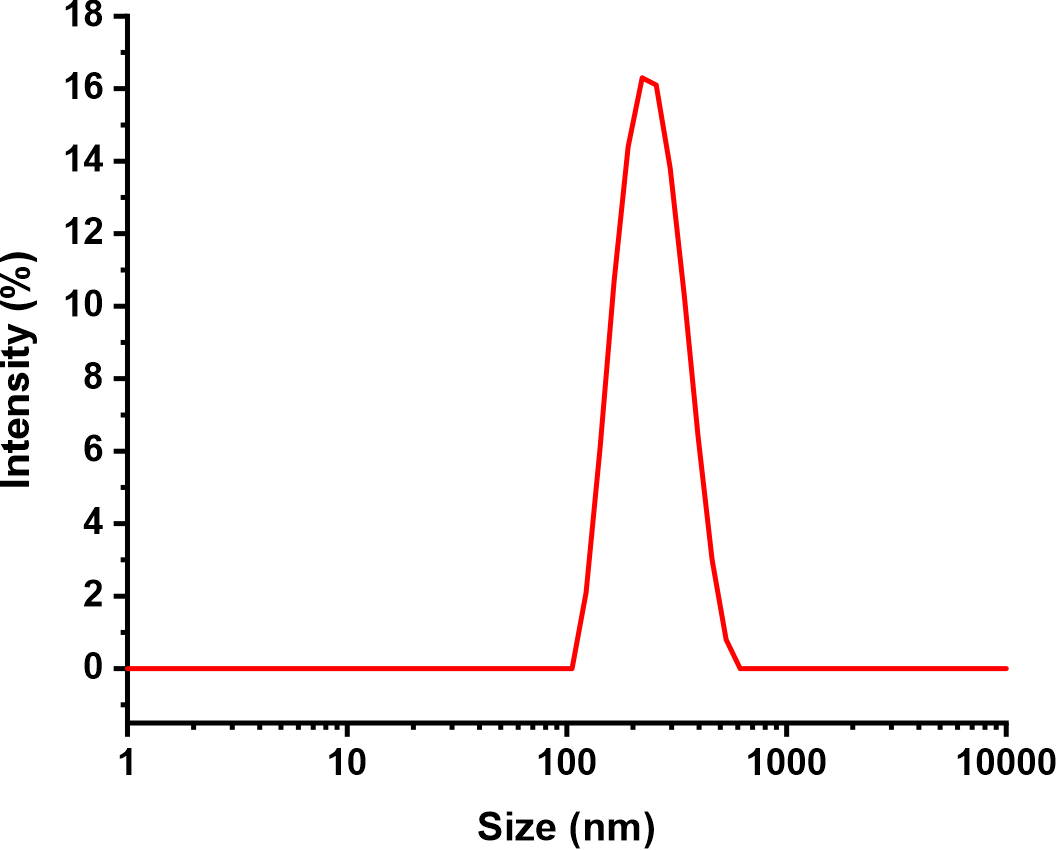

**Figure S1.** Intensity (%) distribution of PPS nanoparticle hydrodynamic size, showing a Z-average size of about 147 nm and a PDI of 0.091

|  | Untreated | | | | | | PPS-NP Treated | | | | | |
| --- | --- | --- | --- | --- | --- | --- | --- | --- | --- | --- | --- | --- |
| Interstitial fibrosis | 0 | 0 | 0 | 0 | 0 | 0 | 0 | 0 | 0 | 0 | 0 | 0 |
| Arteriolar Hyalinosis | 0 | 0 | 0 | 0 | 0 | 0 | 0 | 0 | 0 | 0 | 0 | 0 |
| Vascular Narrowing | 0 | 0 | 0 | 0 | 0 | 0 | 0 | 0 | 0 | 0 | 0 | 0 |
| Acute Tubular Injury | 2 | 2 | 1 | 1 | 1 | 3 | 0 | 0 | 2 | 1 | 2 | 2 |
| Tubular Atrophy | 0 | 0 | 0 | 0 | 0 | 0 | 0 | 0 | 0 | 0 | 0 | 0 |
| Globally Sclerosed Glomeruli | 0 | 0 | 0 | 0 | 0 | 0 | 0 | 0 | 0 | 0 | 0 | 0 |

**Supplementary Table 1 – Histology grading.** Kidney biopsy samples were fixed, sectioned and H&E stained before being graded according to the Remuzzi scoring system by a clinical histopathologist who was blinded to the study groups. Treated kidneys exhibited better preservation of tissue architecture, with less overall acute tubular necrosis.
